# BMI-1 Modulation and Trafficking During M Phase in Diffuse Intrinsic Pontine Glioma

**DOI:** 10.1101/2025.05.16.654605

**Authors:** Banlanjo Umaru, Deepak Kumar Mishra, Shiva Senthil Kumar, Hao-Han Pang, Brendan Devine, Komal Khan, Rachid Drissi

**Author notes:** Corresponding Author: Rachid Drissi, Center for Childhood Cancer Research, Nationwide Children’s Hospital, 575 Children’s Crossroad, Columbus, OH, USA. Contributed equally to this work (Banlanjo Umaru, Deepak Kumar Mishra).

## Abstract

BMI-1 (B cell-specific Moloney murine leukemia virus integration site 1) has been implicated in both normal and cancer cell biology. While the canonical function of BMI-1 involves epigenetic repression, novel extra-nuclear functions have been recently reported. In the present study, we demonstrate that the phosphorylation of BMI-1 in diffuse intrinsic pontine glioma (DIPG) cells occurs in M phase and that triggers simultaneous translocation of the phosphorylated BMI-1 to the cytoplasm. This translocation is mediated by the RanGTP-dependent transporter CRM1, also known as exportin. Furthermore, we uncovered a previously unidentified nuclear export signal (NES) in BMI-1 protein, suggesting an active transport type of modified BMI-1 mediated by CRM1. These findings associate BMI-1 phosphorylation with its trafficking in M phase. Collectively, this study sheds light on the molecular mechanisms underlying BMI-1 functions in DIPG, thereby potentially paving the way for the development of targeted therapeutic strategies related to M phase progression.

## 1. Introduction

BMI-1 is a highly conserved protein initially identified as a proto-oncogene that cooperates with c-MYC in the tumorigenesis of murine B-cell lymphoma [1]. It is a subunit of the Polycomb repressive complex 1 (PRC1) required for the canonical RING1B mediated E3-ubiquitin ligase activity that catalyzes the ubiquitination of histone H2A at lysine 119 (H2A-K119Ub). BMI-1- associated E3 ubiquitin ligase activity represses multiple gene loci, including *INK4A/ARF* locus encoding for two tumor suppressors p16^INK4A^ and p14^ARF^ [2]. BMI-1 has been implicated in a number of biological functions including development, cell cycle, DNA damage response (DDR), senescence, stem cell self-renewal and differentiation [3]. Furthermore, BMI-1 is highly expressed in multiple malignancies and it’s involved in therapy resistance, tumorigenesis, metastasis, and cancer recurrence[3, 4]. These functions are attributed to three domains of BMI-1; the N-terminal RING finger (RF) and the central helix-turn-helix (HTH) domains that are responsible for its interaction with DNA thereby facilitating gene repression, DDR and senescence prevention [5–8]. The C-terminal PEST domain harbors several phosphorylation sites and confers protein stability and increased oncogenic potential in multiple cancers[9–12]. Additionally, BMI-1 contains two nuclear localization sequences (NLS1 and NLS2), with only NLS2 shown to be responsible for its nuclear localization[13, 14].

BMI-1 is tightly regulated both at the transcriptional and the post-translational level. While transcriptional regulation has been extensively studied, the role of post-translational modifications in modulating BMI-1 function is poorly understood[15]. Recent reports have elucidated novel non-nuclear functions of BMI-1. For example, BMI-1 has been shown to localize to the inner mitochondrial membrane supporting the stability of mitochondrial RNA transcripts and thereby regulates mitochondrial function[16, 14]. BMI-1 has also been shown to regulate the expression of androgen receptor in prostate cancer, through direct binding in a PRC1-independent manner[17, 18]. In support of these extra-nuclear functions, BMI-1 has been previously shown to be associated with chromatin when hypo-phosphorylated in G1/S, and dissociates from the chromatin in M phase when phosphorylated[19]. Recently, we showed cytoplasmic translocation of phosphorylated BMI-1 in M phase [20]. However, the process of BMI-1 translocation from the nucleus to the cytoplasm and its role during M phase remain largely unknown. In the current study, we show that the cytoplasmic translocation of BMI-1 is M phase specific and PRC1- independent. This translocation is an active process mediated by exportin (CRM-1). Additionally, we have identified a functional novel nuclear export signal (NES) sequence within the BMI-1 protein that allows transport to the cytoplasm by CRM1.

## 2. Materials and methods

### 2.1. Cell lines

The primary patient-derived DIPG cells SU-DIPG-IV and CCHMC-DIPG-1 were cultured as previously described [20]. SU-DIPG-IV was authenticated by our collaborator M. Monje (Stanford University). CCHMC-DIPG-1 cells were generated and authenticated by the Drissi laboratory using whole genome sequencing (WGS) and RNA-seq. Using the Universal Mycoplasma Detection Kit (ATCC; 30-1012K), all cell lines used in this study were confirmed to be negative for mycoplasma contamination. HEK293T cells (ATCC, RRID: CVCL_0063) were purchased from ATCC and used for lentiviral vector packaging.

### 2.2. Reagents

Thymidine (Sigma, CAS# 50-89-5), Colchicine (Sigma CAS# 64-86-8), Leptomycin-B (Sigma CAS# 87081-35-4) and Nocodazole (Sigma CAS# 31430-18-9) was reconstituted in sterile water. PTC596 was provided by PTC Therapeutics (South Plainfield, NJ, USA) and was reconstituted in DMSO for *in vitro* studies.

### 2.3. Generation of ΔNES-BMI-1 mutants

The NES deletion (ΔNES-BMI-1) cDNA was generated through overlap extension PCR. Wild type (WT) and the NES deletion (ΔNES) BMI-1 cDNA were cloned into pLVX-AcGFP1-N1 Vector (Clontech, #632154) to create GFP-tagged expression vectors. These vectors were transfected into HEK293T cells with additional helper plasmids pMDLg/pRRE (#12251), pMD2.G (#12259), pRSV-Rev (#12253) from Addgene. Supernatant containing viruses were collected 48 hours post-transfection, concentrated and directly added to CCHMC-DIPG-1 cells. GFP positive cells were sorted and expanded. NES deletion was confirmed through Sanger sequencing.

### 2.4. Cell Cycle, Proliferation and Apoptosis Assays

Cell cycle was analyzed as previously described[20]. Data were analyzed using FlowJo v.10 (FlowJo, USA) software. Cell proliferation was measured using WST-1 assay (Takara Bio, MK400) as per manufacturer’s instructions. WST-1 reagent was added to each well at a final concentration of 1:10, incubated for 1.5 hour at 37°C, and absorbance was measured at 450 nm with 650 nm as the reference wavelength.

### 2.5. Western Blotting

Immunoblot assays were performed as previously described[20]. Antibodies used were against BMI-1 (Cell signaling, Cat# 5856, RRID: AB_10838137, 1:1000), H2AK119Ub (Cell signaling, Cat# 8240S, RRID: AB_10891618, 1:1000), H3 S10-P (Cell signaling, Cat# 9706, RRID: AB_331748, 1:1000), Cyclin B1 (Cell signaling, Cat# 4138, RRID: AB_2072132, 1:1000), Cleaved caspase-3 (Cell signaling Cat# 9661, RRID: AB_2341188, 1:1000), Total Histon H3 (Cell signaling Cat# 3638S, RRID: AB_1642229, 1:1000), Total Histon H2A (Cell signaling Cat# 3636, RRID: AB_2118801, 1:1000), β-Actin (Cell signaling Cat# 3700, RRID: AB_2242334, 1:1000).

Membranes were stained with corresponding secondary antibodies, Goat anti-mouse IgG (H+L) Secondary Antibody, HRP (Thermo Fisher Scientific, Cat# 31430, RRID: AB_228307, 1:1000) or Goat anti-Rabbit IgG (H+L) Secondary Antibody, HRP (Thermo Fisher Scientific, Cat# 31460, RRID: AB_228341, 1:1000). Bands visualized with ECL were captured using Azurec500 imaging system (Azure Biosystems) and quantified using Image Studio Lite (LI-COR). For the λ- Phosphatase experiments, 15 µg of protein lysates were treated with λ-Phosphatase (NEB, Cat# P0753S) according to manufacturer’s protocol and then immunoblot was performed. Cytoplasmic and nuclear fractions were isolated as described elsewhere [21].

### 2.6. Immunofluorescence

Immunostaining was performed as described previously [20]. Primary antibodies were used against BMI-1 (Cell signaling, Cat# 6964, RRID: AB_10828713, 1:500), RING1B (Cell signaling, Cat# 5694, RRID: AB_10705604, 1:500), H3 S10-P (Cell signaling, Cat# 9706, RRID: AB_331748, 1:500), Cyclin B1 (Cell signaling, Cat# 4138, RRID: AB_2072132, 1:500) were stained with corresponding secondary antibodies, Alexa Fluor® 488 AffiniPure™ Donkey Anti- Rabbit IgG (H+L) (Jackson ImmunoResearch, Cat# 711-545-152, RRID: AB_2313584, 1:500) or Alexa Fluor® 594 AffiniPure™ Donkey Anti-Mouse IgG (H+L) (Jackson ImmunoResearch, Cat# 715-585-150, RRID: AB_2340854, 1:500). For nuclear staining, cells were embedded with mounting media with DAPI (Vector Laboratories, H1200). Images were captured with 60 X oil objective on Nikon Eclipse Ti confocal microscope. Quantifications were performed using ImageJ software (NIH image).

### 2.7. Statistical analysis

Data from at least two independent experiments with individual technical replicates wherever applicable were collected. Representative images or blots are shown. Results are shown as mean ± SD. GraphPad Prism 8.0.1 was used to perform statistical analysis. One- or Two-way ANOVA followed by a post-hoc Dunnet’s or Tukey test, wherever applicable, was used to analyze the data.

## 3. Results

### 3.1. Phosphorylated BMI-1 is translocated to the cytoplasm in M phase

In our previous studies with PTC596, a mitotic blocker, we showed that treatment with PTC596 led to BMI-1 phosphorylation and this modification correlated with its translocation to the cytoplasm in cells arrested in M phase [20]. To confirm these observations, we synchronized CCHMC-DIPG-1 cells in M phase using nocodazole (Figure 1A) and observed time-dependent accumulation of phosphorylated BMI-1 as the cells arrested in M phase (Figure 1B). Using previously established mitotic blockers, colchicine or PTC596, we observed an accumulation of phosphorylated BMI-1 in the cytoplasmic compartment in cells arrested in M phase (Figures 1C and 1D). Moreover, these observations were further validated in cell synchronization studies using PTC596 (Figures S1A-C). Using another DIPG cell line (SU-DIPG-IV) treated with either PTC596 or colchicine as well as with λ-phosphatase, we showed that modification of BMI-1 occurs via phosphorylation (Figure S1D). Of note, we used Cyclin B1 as a surrogate marker for identifying cells in the G2/M phase in both immunoblot and immunofluorescence (IF) assays. Consistent with previous findings, Cyclin B1 accumulates in the cytoplasm as cells approached prophase[22–24]. Given that BMI-1 is a crucial component of the PCR1 complex, we aimed to investigate whether its phosphorylation and translocation influence the trafficking of the entire complex. Interestingly, RING1B, the catalytic subunit of PRC1 complex, was not translocated to the cytoplasm when treated with PTC596 or colchicine (Figure 1E). Furthermore, IF analysis validated these findings by confirming cytoplasmic translocation of BMI-1 and not of RING1B (Figures 1F and 1G) in M phase cells as evidenced by H3 S10-P positive staining. Together these data suggest that BMI-1 phosphorylation and translocation to the cytoplasm in M phase is PRC1-independent.

**Figure 1.**
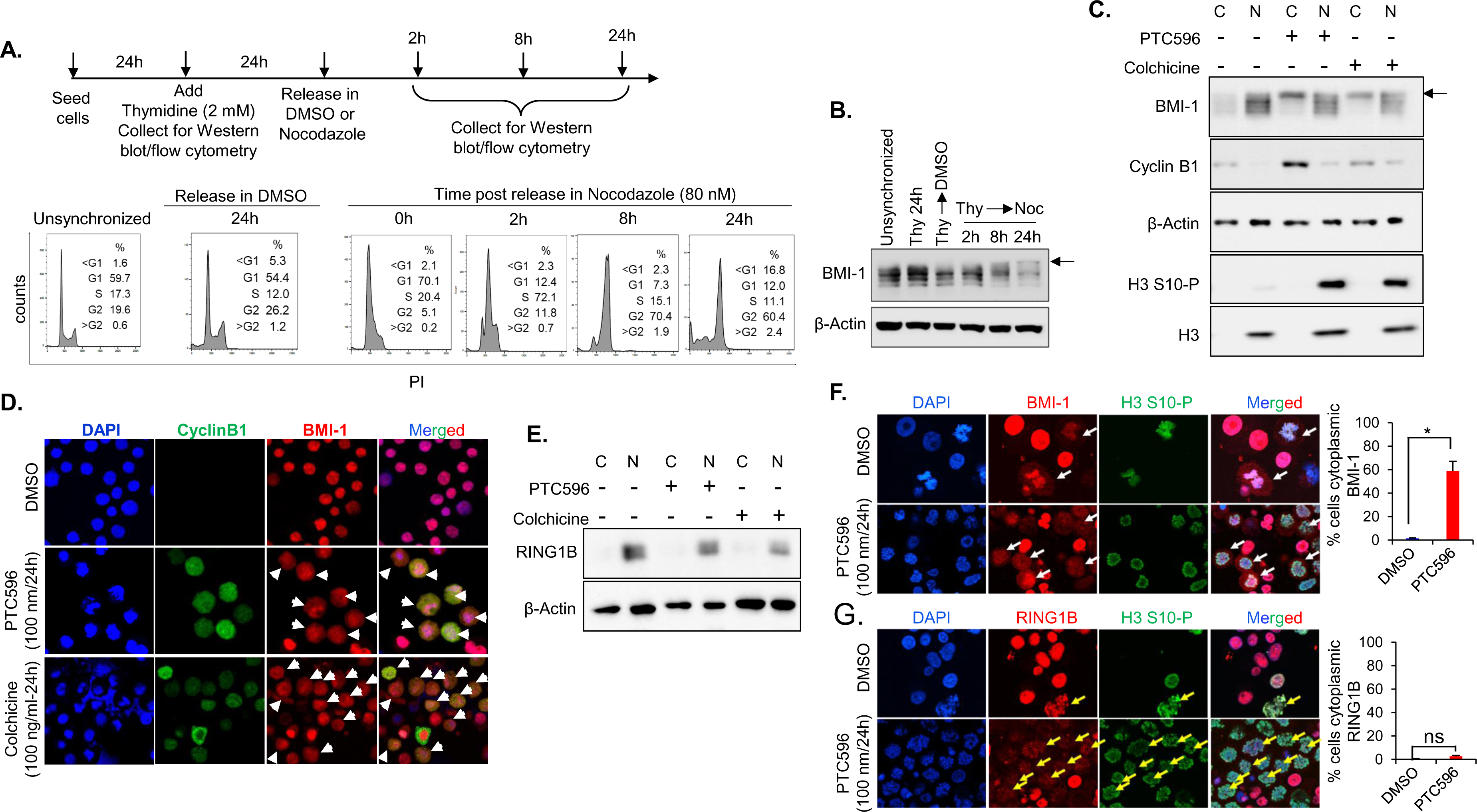
Phosphorylation and translocation of BMI-1 occurs in M phase. **(A)** Scheme for the cell synchronization studies (top) and corresponding cell cycle analysis by flow cytometry (bottom). The percentage of cells in each cell cycle is indicated. (**B)** Immunoblot analysis of BMI- 1 at the indicated timepoints. The arrow indicates phosphorylated BMI-1. β-Actin is used as the loading control. (**C)** Immunoblot analysis of BMI-1, Cyclin B1 and histone H3 S10-P in the cytoplasmic (C) and nuclear (N) fractions when treated with PTC596 (100 nM for 24 h) or colchicine (100 ng/ml for 24 h). β-Actin and total H3 served as loading controls. Arrow indicates phosphorylated BMI-1. (**D)** Representative immunofluorescence images showing DAPI (blue), Cyclin B1 (green) and BMI-1 (red) in cells treated with PTC596 or colchicine at the indicated doses for 24 h. White arrows indicate the cells with cytoplasmic localization of BMI-1. (**E)** immunoblot analysis of RING1B in the cytoplasmic (C) and nuclear (N) fractions in cells treated with PTC596 (100 nM for 24h) or colchicine (100 ng/ml for 24h). (**F)** Representative immunofluorescence images showing DAPI (blue), H3-S10-P (green) and RING1B (red) in cells treated with 100 nM PTC596 for 24 h. White arrows indicate the cells with cytoplasmic localization of BMI-1. **(G)** Representative immunofluorescence images showing DAPI (blue), H3-S10-P (green) and RING1B (red) in cells treated with 100 nM PTC596 for 24 h. Yellow arrows indicate the cells with RING1B still localized within DNA. All experiments were conducted in CCHMC- DIPG-1 cells at n=2. P values are indicated (*, P < 0.05; ns, not significant (P>0.05))

### 3.2. Transport of modified BMI-1 to the cytoplasm is CRM1 dependent

When treated with PTC596, we observed a small percentage of cells showing both cytoplasmic and nuclear BMI-1 distribution (Figure S2A), suggesting that BMI-1 may be actively transported from the nucleus to the cytoplasm during M phase. Nuclear export of majority of proteins to the cytoplasm are directed by the nuclear export receptor CRM1 or exportin-1[25, 26]. Leptomycin-B (LMB), a potent selective inhibitor of CRM1-mediated transport, has been previously reported to inhibit nuclear export of Cyclin B1 in a time-dependent manner[27, 24]. We observed a gradual decrease in cytoplasmic BMI-1 and a consequent increase in the nuclear levels in a time-dependent manner when treated with LMB (Figure 2A). Interestingly, when treated together with PTC596, LMB inhibited the cytoplasmic translocation of phosphorylated BMI-1 in M phase (Figures 2B and D). Moreover, LMB treatment interfered with PTC596-induced M phase arrest (Figure 2C). Together, these data indicate that the BMI-1 translocation to the cytoplasm in M phase is an active process and is CRM1-dependent.

**Figure 2.**
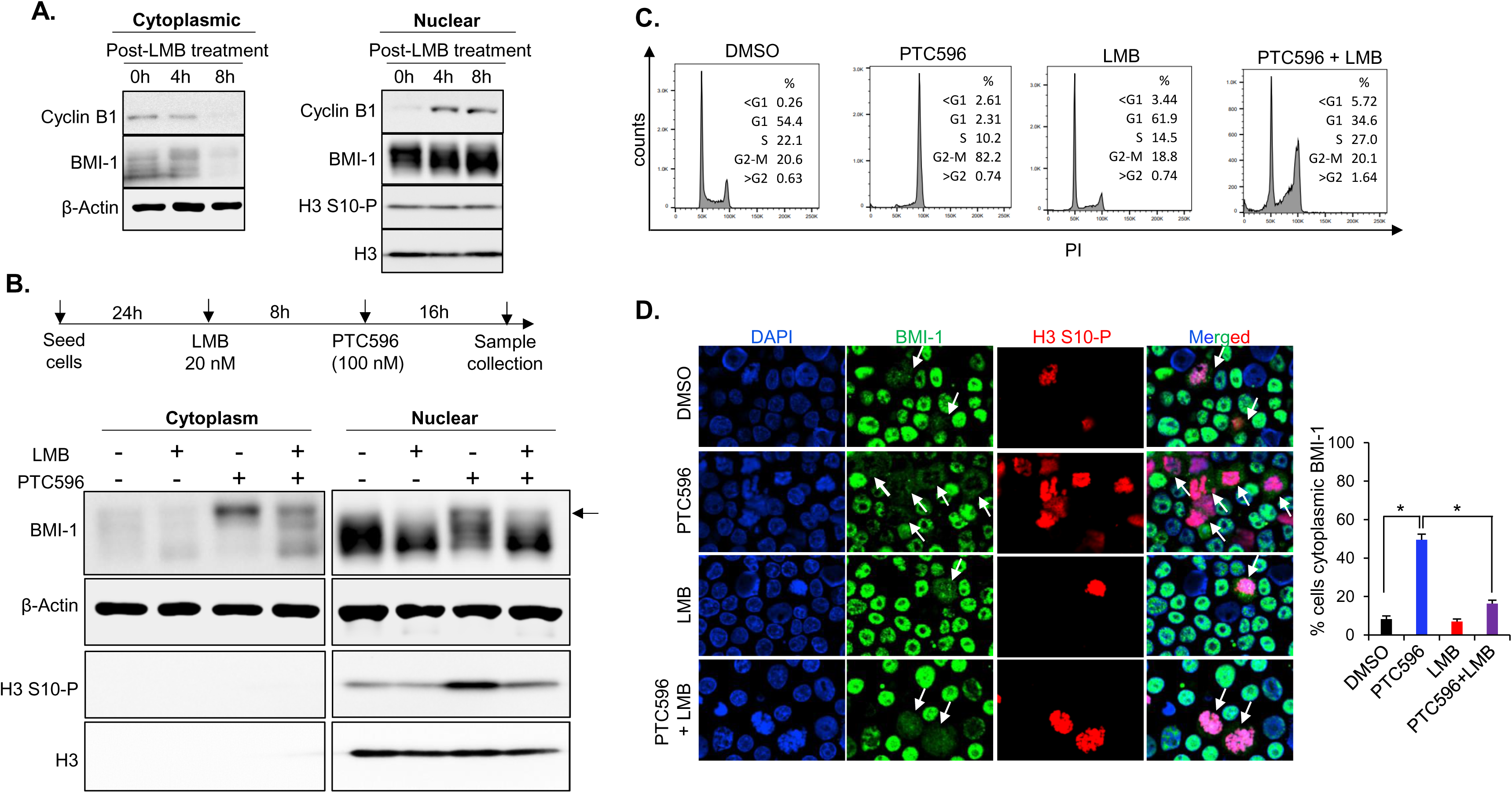
M phase translocation of BMI-1 is an active process. **(A)** Immunoblot analysis of BMI-1, Cyclin B1 and H3 S10-P in the cytoplasmic and nuclear fractions of cells treated with leptomycin B (LMB) for the indicated time points. β-Actin and total H3 served as the loading controls. **(B)** Scheme for the combination experiment of LMB with PTC596 (top) and corresponding immunoblot analysis of BMI-1 and H3 S10-P in the cytoplasmic and nuclear fractions (bottom). β-Actin and total H3 served as loading controls. (**C)** Cell cycle analysis by flow cytometry of cells treated with LMB and PTC596. The percentage of cells in each cell cycle phase is indicated. **(D)** Representative immunofluorescence images showing DAPI (blue), BMI-1 (green) and H3 S10-P (red) in cells treated with PTC596 and LMB based on scheme in **(B).** White arrows indicate the cells with cytoplasmic localization of BMI-1. All experiments were conducted in CCHMC- DIPG-1 cells at n=2. P values are indicated (*, P < 0.05)

### 3.3. Characterization of functional NES motif in BMI-1

CRM1 mediated transport requires a leucine-rich sequence in the target cargo proteins that have been characterized as nuclear export signals (NES)[28, 29]. Using an NES prediction pipeline [30], we identified a previously unidentified putative NES sequence in BMI-1 protein, 175- LRKFLRSKMDI-185 within HTH domain (Figure S3A). This sequence conforms to the NES consensus Class 1C sequence (φ-XXX-φ-XXX-φ-X-φ)[31, 32] and is highly conserved across multiple species (Figures 3A and B). Additionally, mutations in the *CRM1* gene has been previously shown to interfere with the transport function of the CRM1 protein[33]. We confirmed that CCHMC-DIPG-1 cells do not harbor any mutations in the *CRM1* gene. To examine whether the putative NES sequence is functional, we ectopically expressed ΔNES-BMI-1 (D1- ΔNES-BMI- 1) as well as the BMI-1 overexpression control (D1-BMI-1) in CCHMC-DIPG-1 cells (Figure S3B). When treated with PTC596, we observed both a decrease in phosphorylation in D1-ΔNES-BMI-1, and a defect in phosphorylated BMI-1 cytoplasmic translocation compared to D1-BMI-1. Importantly, the ectopic expression of ΔNES-BMI-1 did not affect the phosphorylation and cytoplasmic translocation of endogenous BMI-1 (Figures 3C and S3A). Moreover, the ectopic expression of ΔNES-BMI-1 did not affect the M phase arrest of cells treated with PTC596 (Figures 3C and D), suggesting that a novel NES sequence identified in BMI-1 may be specifically involved in conformational change and translocation of phosphorylated BMI-1 in M phase.

**Figure 3.**
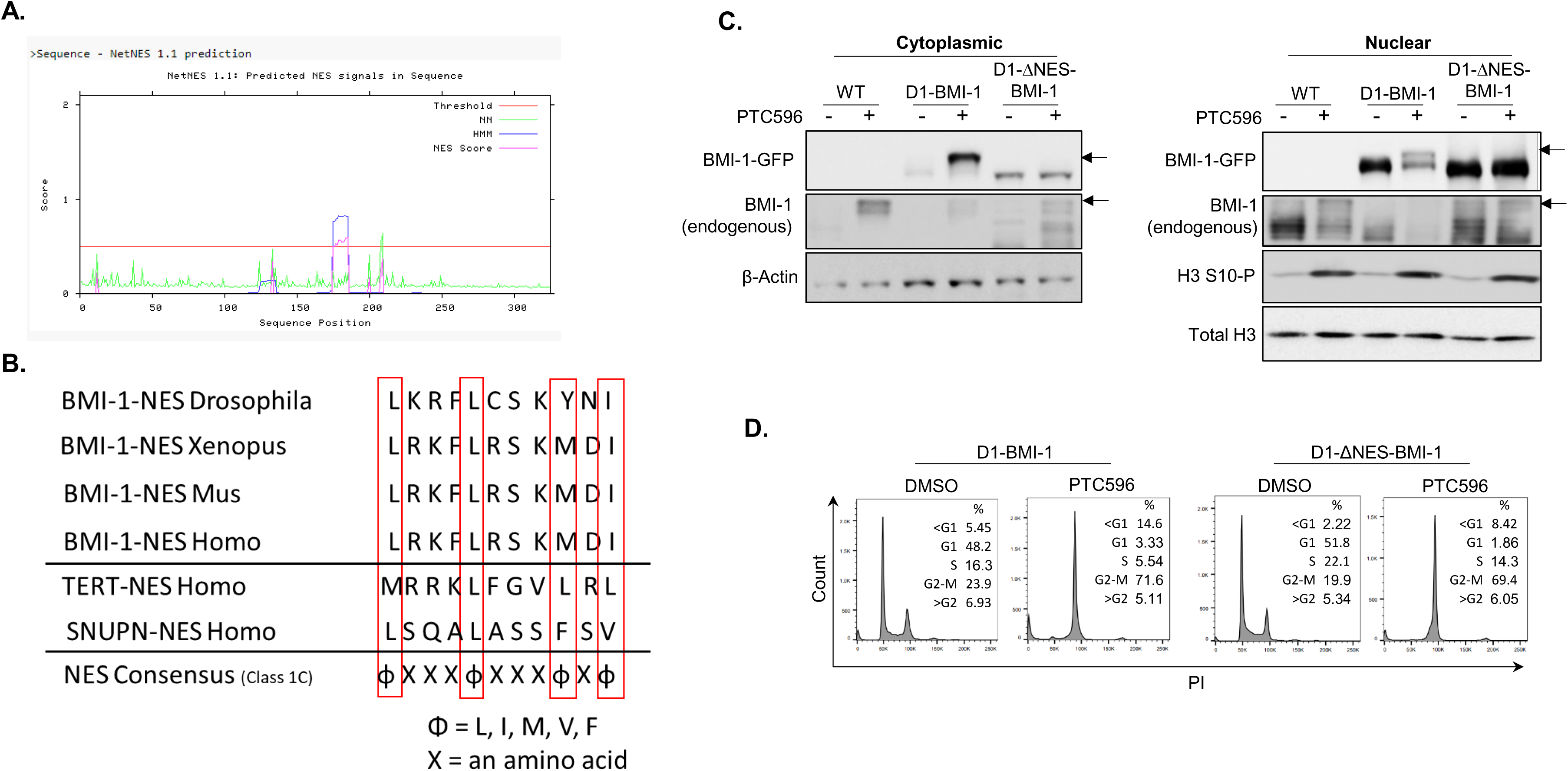
Identification and characterization of NES in BMI-1. **(A)** Output of the NES prediction pipeline NetNES 1.1 showing putative NES signals in the BMI-1 protein sequence (NN = neural network algorithm; HMM = hidden Markov Model algorithm; NES score = combination of NN and HMM algorithms). A signal above the threshold signifies a highly probable NES hit. **(B)** Alignment of the identified BMI-1 NES sequence in multiple species, verified NES sequences in TERT and SNUPN and the NES consensus sequences showing high homology. **(C)** Immunoblot analysis of BMI-1, H3 S10-P from cytoplasmic (C) and nuclear (N) fractions of D1-BMI-1 and D1-ΔNES-BMI- 1 cells treated with PTC596. β-Actin and total H3 served as loading controls. BMI-1-GFP is ectopically expressed in D1-BMI-1 and D1-ΔNES-BMI-1. Arrows indicate phosphorylated BMI-1. **(D)** Cell cycle analysis by flow cytometry of cells of D1-WT-BMI-1 and D1-ΔNES-BMI-1 cells treated with DMSO or PTC596 (100 nM for 24h). The percentage of cells in each cell cycle phase is indicated.

## 4. Discussion

Diffuse Intrinsic Pontine Glioma (DIPG) represents one of the most aggressive malignancies of the central nervous system, predominantly affecting children with a median overall survival of less than one year. Hence, there is an urgent need to develop novel therapies that not only improve outcome but mitigate long-term complications in children with DIPG. We have previously identified BMI-1 as a potential therapeutic target in DIPG and have shown that BMI-1 is highly expressed in DIPG tumors regardless of DIPG subtype[34]. The protein BMI-1, a critical component of the Polycomb Repressive Complex 1 (PRC1), is a proto-oncogene implicated in development, stemness of normal and malignant cells, and self-renewal. Elucidating the molecular mechanisms underlying BMI-1 functions in DIPG-M phase progression will contribute to the development of targeted therapeutic strategies.

We have previously reported the phosphorylation of BMI-1 in M phase and the simultaneous translocation to the cytoplasm of modified BMI-1 when DIPG cells were treated with PTC596[20]. Our present study demonstrates that the phosphorylation and translocation of BMI-1 is independent of the PRC1 complex. Interestingly, the translocation is an active process mediated by CRM1, an essential mediator of nuclear protein export. Voncken *et al.* using osteosarcoma cells, showed that BMI-1 was phosphorylated at the G2/M phase of the cell cycle and dissociates with PRC1 complex proteins from chromatin[35, 36]. However, in this study, we observed colocalization of RING1B, the catalytic subunit of the PRC1 complex, with chromatin and no cytoplasmic translocation during M phase in our DIPG cells treated either with colchicine or PTC596 (Figures 1E and 1F). This suggests that the phosphorylation and translocation to the cytoplasm of BMI-1 is PRC1-independent and indicate a non-canonical role of BMI-1 in DIPG cells. Moreover, we have previously demonstrated high expression levels of BMI-1 and normal levels of RING1B in DIPG tumor tissues in all subtypes of patient-derived DIPG neurospheres[34], suggesting extranuclear PRC1-independent and non-canonical functions of BMI-1. Further studies are required to elucidate these extranuclear functions of phosphorylated BMI-1 in DIPG. Extranuclear BMI-1 has been shown to localize to the inner mitochondrial membrane to stabilize the mitochondrial RNA, thereby regulating the electron transport chain, which enhances oxidative phosphorylation and ATP synthesis and prevents aberrant ROS production[37, 38].

During mitosis, the RanGTPases, including CRM1, control multiple cellular processes involving nuclear transport, mitotic checkpoints, spindle assembly, and post-mitotic nuclear envelope reassembly[25, 26]. Here, we show that phosphorylated BMI-1 is transported actively from the nucleus to the cytoplasm via CRM1. A comprehensive analysis of the mitotic events is required to determine the timing of BMI-1 phosphorylation and consequent transport from the nucleus to the cytoplasm during M phase. We have previously postulated that BMI-1 phosphorylation occurs in early metaphase after induction of the spindle assembly checkpoint (SAC) and before the anaphase promoting complex (APC/C^CDC20^) inhibition[20], albeit Kim *et al.* have suggested that PTC596 directly inhibits the APC/C^CDC20^ leading to persistent cyclin-dependent kinase (CDK)1/2 activity and BMI-1 hyper-phosphorylation, as well as reduced PRC1 activity[39]. Moreover, we have also shown that BMI-1 phosphorylation coincides with DNA damage and apoptosis in DIPG cells but not in normal cells, suggesting a survival role of BMI-1 in M phase in DIPG cells involving DDR[20].

In this study, we identified for the first time an NES sequence in BMI-1 protein involved in its CRM1-mediated nuclear export. Indeed, previous studies have reported an NES-dependent nuclear export of Cyclin B1 by CRM1[22–24]. Moreover, treatment with LMB, which was shown to inhibit interactions of CRM1 with NES[40], coupled with Cyclin B1 accumulation in the nucleus led to cell cycle arrest in S and G2[24]. Furthermore, LMB inhibition of CRM1 was shown to disrupt mitotic progression and chromosome segregation[41]. As expected, we observed an inhibition of PTC596 induced M phase arrest when cells were treated with LMB thus interfering with modified BMI-1 translocation.

Several questions remain to be answered to elucidate the exact role of BMI-1 phosphorylation in tumor survival and progression. We are currently performing further studies to determine the phosphorylation sites, upstream kinases, effects of phosphorylation on structural modification of BMI-1, new binding partners of modified BMI-1, and ultimately the extranuclear locations and non-canonical functions of BMI-1.

## 5. Conclusion

In conclusion, our findings indicate a potential non-canonical function of BMI-1 in DIPG during the M phase. BMI-1 undergoes phosphorylation during M phase, dissociates from the PRC1 complex, and subsequently actively translocates to the cytoplasm (Figure 4). Future studies including site directed mutagenesis of Lysine residues in the NES domain should elucidate its role in BMI-1 transport. Additionally, exploring and targeting the BMI-1 phosphorylation sites is prerequisite to uncover the extranuclear locations and functions of phosphorylated BMI-1 in DIPG.

**Figure 4.**
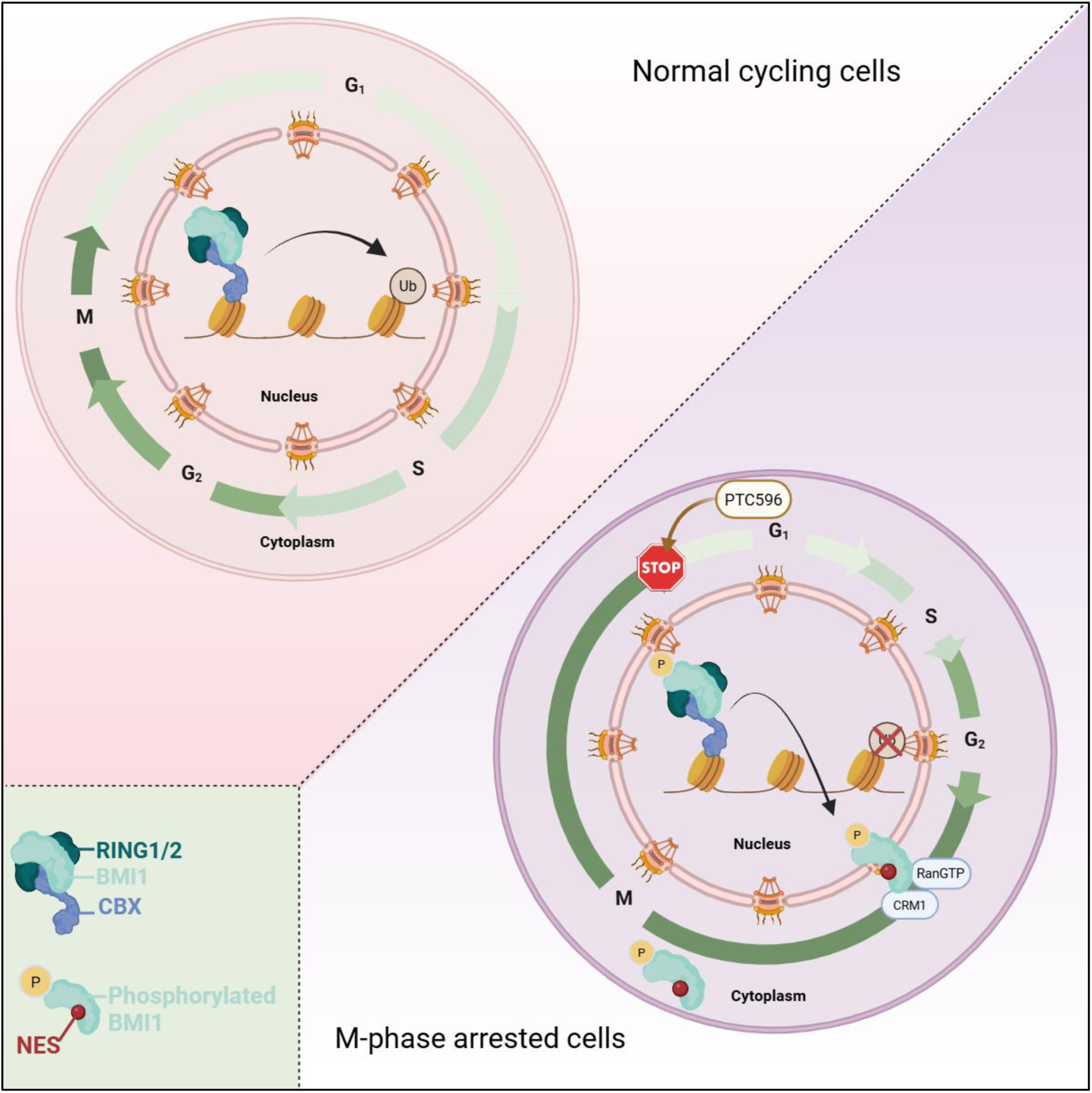
Identification and characterization of NES in BMI-1. Schematic illustration of BMI-1 active translocation from the nucleus to the cytoplasm following phosphorylation. In normal cycling cells, BMI-1 in the PRC1 complex, facilitates the monoubiquitination of histone H2A at lysine 119. However, in M phase arrested cells induced by PTC596, BMI-1 is phosphorylated leading to its dissociation from the PRC1 complex, and subsequent translocation to the cytoplasm mediated by its NES and CRM1.

## Acknowledgements

This work was supported by the Department of Defense grant 13709017, the National Center for Advancing Translational Sciences grant UM1TR004548, and Nationwide Children’s Hospital, Columbus, OH. We thank M. Monje (Stanford University) for kindly providing us with SU-DIPG-IV cells. We thank PTC Therapeutics, Inc. for providing PTC596.

## Author contributions

Conception and design: D.K.M, B.U., S.S.K., H-H.P., and R.D.

Development of methodology: D.K.M., S.S.K., B.U., H-H.P., and R.D.

Acquisition of data: D.K.M., S.S.K., B.D., and U.B.

Analysis and interpretation of data: D.K.M., S.S.K., B.U., and R.D. Writing, review, and/or revision of the manuscript: all authors.

Administrative, technical, or material support: B.D., and K.K. Study supervision: R.D.

## Data and code availability

Data was analyzed using FlowJo v.10, Image Studio Lite, GraphPad Prism 10, and NIH Image

1. J. This paper does not report original code. Scripts and code utilized in this study are available from the lead contact upon request.

## Supplemental information

Additional supporting information Figures S1–S3 is attached.

## Abbreviations

BMI-1: B cell-specific Moloney murine leukemia virus integration site 1
DIPG: diffuse intrinsic pontine glioma
CRM1: Chromosomal Region Maintenance 1
NES: nuclear export signal
PRC1: Polycomb repressive complex 1
DDR: DNA damage response
LMB: Leptomycin-B HTH: helix-turn-helix

---

**Figure S1.**
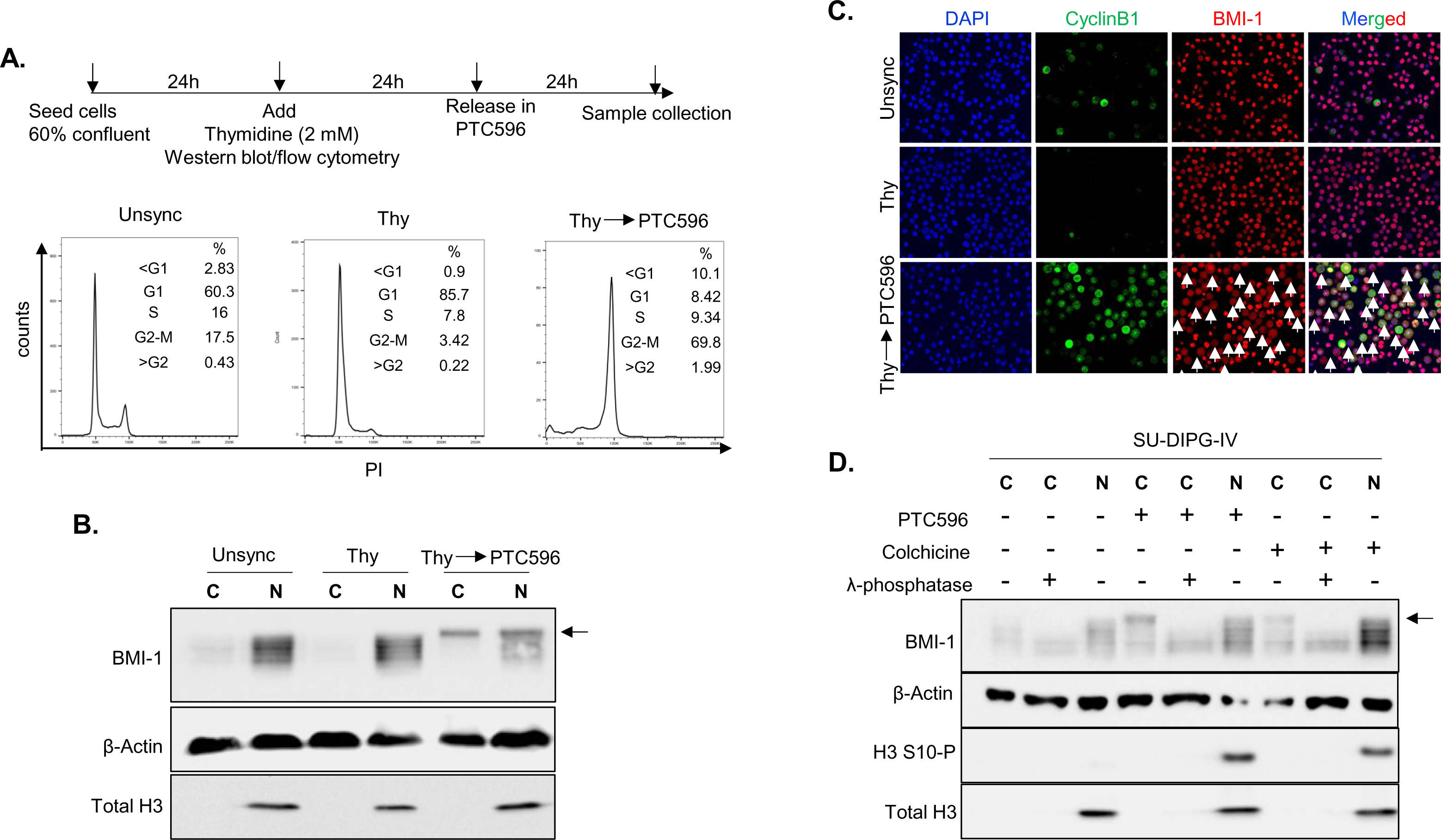
**(A)** Scheme representing the cell synchronization studies with thymidine block and release in PTC596, along with collection timepoints for cell cycle analysis (with indicated percentage of cells). **(B)** Immunoblot analysis of BMI-1 from cytoplasmic (C) and nuclear (N) fractions. Actin and total H3 served as loading control. Arrow indicates phosphorylated BMI-1. **(C)** Representative immunofluorescence images of Cyclin B1 (green) and BMI-1(red). DAPI (blue) represent nuclei. White arrows indicate the cells with cytoplasmic localization of BMI-1. CCHMC-DIPG-1 cells were used for the above experiments. **(D)** Immunoblot analysis of BMI-1 from cytoplasmic (C) and nuclear (N) fractions of SU-DIPG-IV cells treated with PTC596 (100 nM for 24h) or colchicine (100 ng/mL for 24 h). Cytoplasmic fractions were further treated with or without λ-phosphatase. Arrow indicated phosphorylated BMI-1.

**Figure S2.**
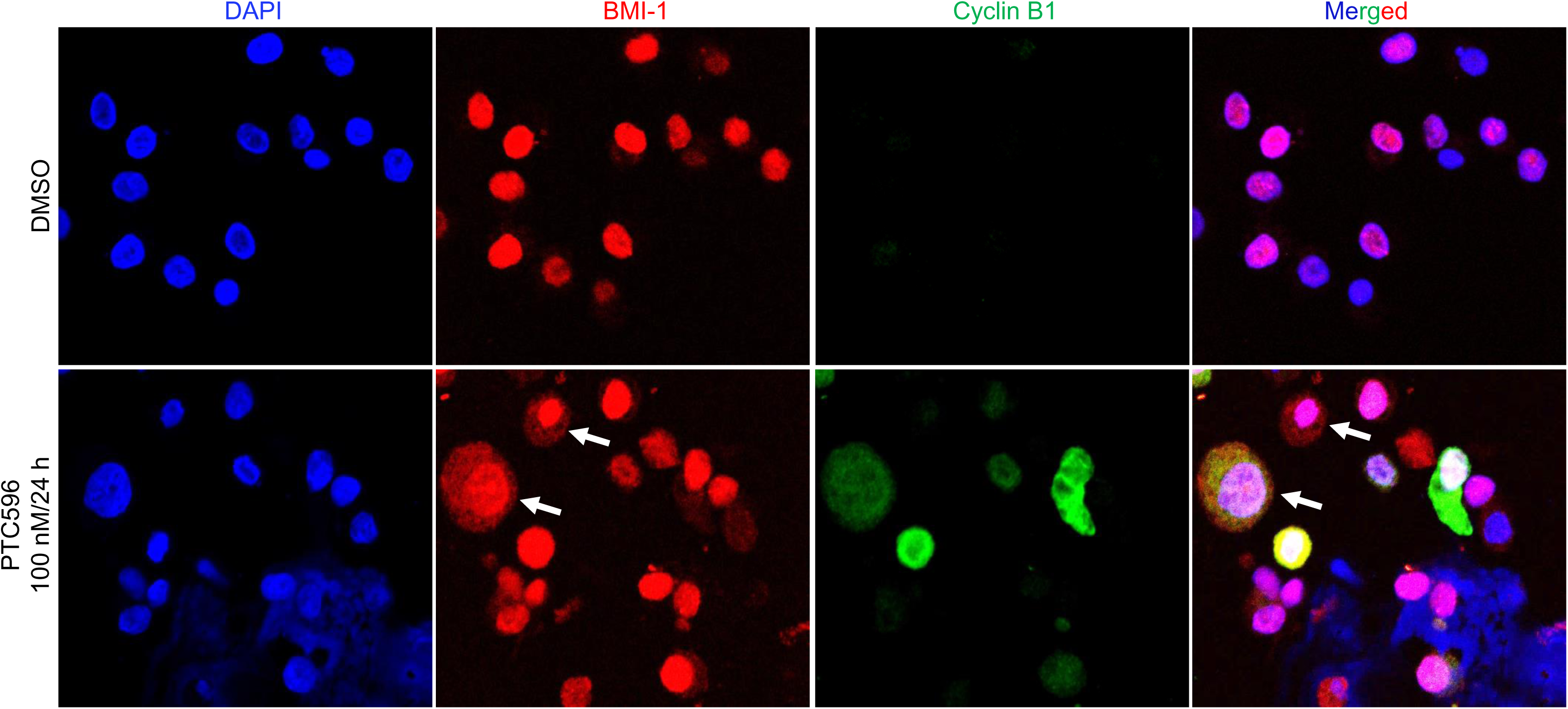
Representative immunofluorescence images of BMI-1 (red) and Cyclin B1(green). White arrows indicate cells with both cytoplasmic and nuclear BMI-1. DAPI (blue) represents nuclei.

**Figure S3.**
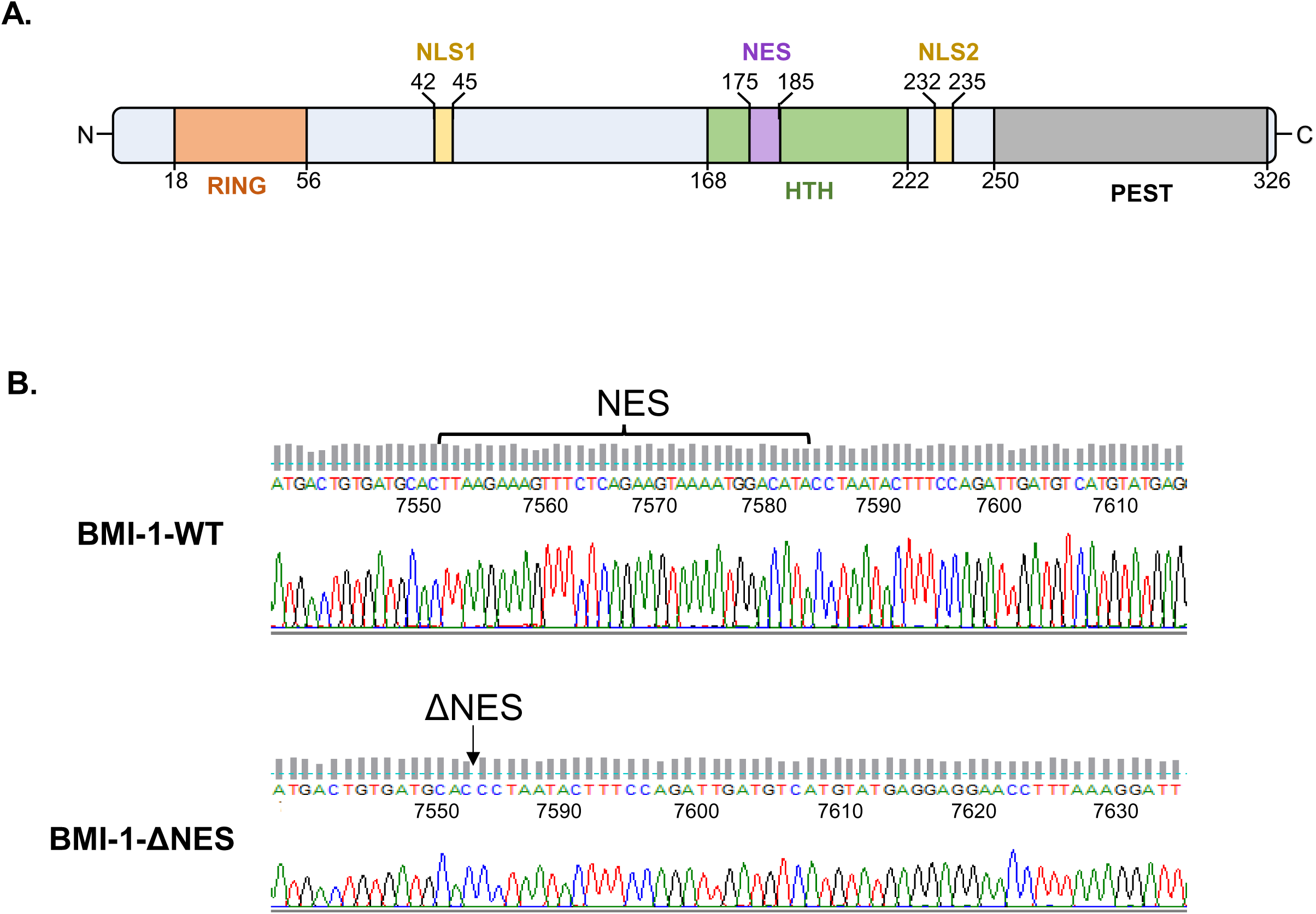
**(A)** Schematic illustration of different domains of the BMI-1 protein structure. The newly identified NES domain within the HTH domain is indicated **(B)** Genomic DNA extracted from D1-BMI-1 and D1-ΔNES-BMI-1 cells were PCR amplified, and Sanger sequenced to confirm deletion of the NES sequence. The numbers indicate the nucleotide position in *BMI-1* gene.

